# The IgCAM CAR regulates gap junction mediated coupling on embryonic cardiomyocytes and affects their beating frequency

**DOI:** 10.1101/2022.11.02.514878

**Authors:** Claudia Matthaeus, René Jüttner, Michael Gotthardt, Fritz G. Rathjen

## Abstract

The IgCAM coxsackie-adenovirus receptor (CAR) is essential for embryonic heart development and for electrical conduction in the mature heart. However, it is not well-understood how CAR exerts these effects at the cellular level. To address this question, we analysed the spontaneous beating of cultured embryonic hearts and cardiomyocytes from wildtype and CAR knockout (KO) embryos. Surprisingly, in the absence of CAR, cultured cardiomyocytes showed an increased frequency of beating and calcium cycling. Increased beating of heart organ cultures was also induced by application of reagents that bind to the extracellular region of CAR, such as the adenovirus fiber knob. However, the calcium cycling machinery, including calcium extrusion via SERCA2 and NCX, was not disrupted in CAR KO cells. In contrast, CAR KO cardiomyocytes displayed an increase in the size, but decrease in total number, of membrane-localized C×43 clusters. This was accompanied by improved cell-cell coupling between CAR KO cells, as demonstrated by increased intercellular dye diffusion. Our data indicate that CAR may modulate the localization and oligomerization of C×43 at the plasma membrane, which could in turn influence electrical propagation between cardiomyocytes via gap junctions.

**Simple summary:** CAR is implicated in higher-order gap junction structure by regulating clustering of connexins in embryonic cardiomyocytes. In the absence of CAR, cardiomyocytes in culture showed an increased frequency of beating and calcium cycling. Perturbation experiments suggest that CAR could be a potential new candidate in cardiac diseases.

## 1. Introduction

The coxsackie-adenovirus receptor (CAR) is a 46 kDa cell adhesion protein of the Ig superfamily. It is composed of the membrane-distal V-type domain (D1) and the membrane-proximal C2-type domain (D2), which are connected by a short junction (Matthaus et al., 2017). CAR shares sequence homology with the IgCAMs CLMP (<u>C</u>AR-like <u>m</u>embrane protein, also called ACAM, adipocyte adhesion molecule), ESAM (<u>e</u>ndothelial cell-<u>s</u>elective <u>a</u>dhesion <u>m</u>olecule) and BT-IgSF (<u>b</u>rain-<u>t</u>estis <u>IgSF</u>, also called IgSF11 or VSIG-3) [4]. During embryonic development, CAR is expressed in various organs, such as the brain, retina, liver, heart, pancreas and lung, and is often found at cell-cell contact sites, including tight junctions [2,5–9]. Shortly after birth, CAR expression significantly decreases in the nervous system and the heart [2,6,10–12]. In the mature heart, CAR specifically localizes to intercalated discs in close association with connexin 43 (C×43), connexin 45 (C×45) and Zona occludens 1 (ZO-1) [13–17].

The absence of CAR in mice causes malformation of the developing heart and hemorrhage, which leads to embryonic lethality around E11.5 to E13.5 [10,13,18]. Conditional inactivation of CAR in the mature heart is associated with reduced expression of C×45 and C×43, which disrupts electrical conduction between the atrium and the ventricle, and leads to atrioventricular block [14,17,19]. In addition, mice with CAR overexpression develop severe cardiomyopathy and die by 4 weeks of age [20]. Interestingly, CAR is expressed at higher levels upon cardiac remodelling in patients suffering from dilated cardiomyopathy [21,22], during myocarditis [23,24] and after myocardial infarction [25]. The normally weak expression of CAR in healthy adult cardiomyocytes, in contrast with its expression in the developing and diseased heart, implicates it in the formation of a functional myocardium and remodelling after injury [26]. The function of CAR in the developing and adult nervous system, however, is still poorly understood. It is implicated in adult neurogenesis, synaptic homeostasis, synaptic plasticity and/or neurite growth [2,27,28].

Adhesion and binding assays, as well as crystallographic studies, indicate that CAR promotes homophilic and heterophilic binding between neighbouring cells [2,11,29,30]. CAR exhibits heterophilic binding to extracellular matrix glycoproteins and to other IgCAMs such as JAML and JAM-C [2,31–33]. The cytoplasmic tail of CAR contains a class I PDZ binding motif for which several binding partners have been identified, such as ZO-1, MUPP-1 (Multi-PDZ domain protein-1), MAGI-1b (Membrane associated guanylate kinase, WW and PDZ domain containing 1b), PICK-1 (Protein interacting with C kinase 1), the synaptic scaffolding protein PSD-95 (postsynaptic density protein 95), LNX (Ligand-of-Numb protein-X) and LNX2 [5,7,34–37].

Previous studies of CAR have focused on its structural aspects and the physiological outcomes when CAR is deleted *in vivo*. However, there is a lack of mechanistic insight into the involvement of CAR in regulating the murine heartbeat. To better understand the molecular and cellular functions of CAR, we analysed its role in the spontaneous beating of embryonic murine hearts and monolayer cardiomyocyte cultures. We show that in the absence of CAR, the beating frequency of embryonic cardiomyocytes is increased. This was correlated with increased calcium cycling and calcium extrusion mechanisms. Further, the increased beating frequency was accompanied by increased gap junction activity, indicated by enhanced dye spread between CAR knockout (CAR KO) cardiomyocytes and an increased size of C×43 clusters. Taken together, our data indicate that CAR is involved in the regulation of C×43 and C×45 localization, which in turn affects the beating of cardiomyocytes at embryonic stages.

## 2. Materials and Methods

### Mice

The mouse line CAR (B6.Cg-Cxadr^tm1^/Fgr) was genotyped as detailed elsewhere [10]. Animals were housed on a 12/12 h light/dark cycle with free access to food. The animal procedures were performed according to the guidelines from directive 2010/63/EU of the European Parliament on the protection of animals used for scientific purposes. All experiments were approved by the local authorities of Berlin (LaGeSO) (numbers T0313/97 and X9014/15).

### Embryonic cardiomyocyte and heart organ cultures

Global CAR-deficient mice die between embryonic day 11.5 and 13.5 [10]. Therefore, organ culture or dissociated cardiomyocytes were prepared from 10.5/11.5 days-old embryos of wild type or CAR mutant mice of the same litter. Intact hearts were cultured in 12-well chamber slides (Ibidi #81201). Hearts were seeded in 50 μl Matrigel (Corning #356230) diluted 1:2 with DMEM/FCS [see also [38]] and incubated for 30 minutes at 37 °C in a humidified atmosphere of 95% air and 5% CO_2_. After solidification of the Matrigel 350 μl of DMEM/FCS with or without of the fiber knob (0.5 mg/ml) or the anti-CAR antibodies Rb80 (final concentration: 0.5 mg/ml of the IgG fraction) to mouse CAR or Rb54 to chick CAR were added. The IgG fractions of Rb80 or Rb54 were obtained by ProteinA (GE healthcare) affinity chromatography. Antibodies were dialyzed against DMEM before application to cultures. Controls of treated cultures contained vehicle and additional control cultures contained the IgG fraction of rabbit antibodies to chick CAR (Rb54) which does not react with mouse CAR [2]. The activity of the fiber knob Ad2 was controlled on CAR-deficient embryonic hearts.

Single cells were obtained by incubating embryonic hearts in 1 mg/ml trypsin in 1mM EDTA/PBS (Gibco) for 10 min at 37^°^C followed by trituration in DMEM supplemented with 10% FCS and penicillin/streptomycin (Gibco). Dissociated cells from one embryonic heart were plated on one glass coverslips (diameter 12 mm) pre-coated with bovine fibronectin (10 μg/ml in PBS; Sigma-Aldrich) in 50 μl DMEM/10% FCS/penicillin/streptomycin at 37 °C in a humidified atmosphere of 95% air and 5% CO_2_. When cells were attached after 4 to 5 hours 450 μl culture medium was added and cells were cultured for 4 to 5 days before measured. Cell or heart organ beatings were counted under an inverted microscope manually at 37°C in a humidified atmosphere with 5% CO_2_. 50% of the medium was exchanged every second day.

### Preparation of the fiber knob of the adenovirus

A cDNA encoding the fiber knob of the adenovirus Ad2C428N (abbreviated Ad2 in the following) was recombinantly expressed in bacteria and purified by sequential steps of ammonium sulphate precipitation, anion exchange chromatography and by Ni-NTA-Agarose (Qiagen #1018244) affinity chromatography as described previously [2,39–41]. Before application to cultures the fiber knob was dialyzed against DMEM.

For whole mount staining of embryonic hearts Ad2 was directly labelled via an NHS ester by the fluorescent dye Cy2 according to the instructions of the manufacturer (GE Healthcare, #32000). Ad2-Cy2 was applied at a concentration of 6 μg/ml in PBS/0.1%Tx100/5% goat serum for 2 hr at room temperature to paraformaldehyde fixed E10.5 hearts followed by several washing steps. Hearts were mounted in Immu-Mount (Thermo Scientific, #9990402) for confocal imaging.

Binding of Ad2 to the fusion proteins CAR-Fc, CLMP-Fc, BT-IgSF-Fc or to the Fc-fragment was done by the ELISA method [42]. 200 μl of 1 μg/ml of the fusion were immobilized on ELISA plates (Nunc) and after blocking of residual binding sites on the plates 200 μl of Ad2 at a concentration of 125 ng/ml was applied. Binding of Ad2 was detected by an HRP-conjugated monoclonal antibody to His-tag (Qiagen) expressed by the fiber knob. O-Phenylenediamine (Sigma) was used as chromogen.

### Calcium imaging of cultured cardiomyocytes

Calcium transients of cultured embryonic cardiomyocytes at DIV4-5 were measured after incubating myocytes with the calcium-sensitive fluorescence dye Fura2-AM (1 μM, Biotium, Cat: 50033-1) diluted in 0.02% Pluronic acid F-127/DMSO (Invitrogen, P3000MP) in culture medium. After incubation for 60 min at 37 °C cells were washed with DMEM and calcium imaging analysis was performed in ACSF (artificial cerebrospinal fluid) (130 mM NaCl, 4 mM KCl, 1.25 mM NaH_2_PO_4_, 25 mM NaHCO_3_, 10 mM Glucose, 2mM CaCl_2_, 1 mM MgCl_2_). Recordings were obtained with Zeiss Axio Examiner A1 and TILL Photonics LA software (version 2.3.0.18, 2013) at room temperature. The ratio (R) of Fura2 fluorescence intensity (R = F_340nm_/F_380nm_) was recorded by excitation at two wavelengths (340 nm and 380 nm, emission wavelength 510 nm, 10-20 ms exposure time, 4×4 binning) was recorded and analysis was carried out with Origin software (Additive, version 7.03, 2002), IgorPro (WaveMetrics, 2014) and Fiji (ImageJ, version 1.49m, 2015). To analyze the rate constants for the different calcium extrusion components (SERCA2, NCX, PMCA and mitochondria) the protocol of Voigt et al. [43] was applied and described in detail in the supplemental information. Blockers were diluted in DMSO and added to the ACSF buffer before recordings were started.

Exposure time of excitation and cycle time depended on Fura-2 loaded cells and the aim of the experiment. For kinetic analysis of calcium transients (frequency, amplitude, decay time constant) the exposure time of 340 nm and 380 nm excitation wavelength was reduced to 10 ms, the cycle time was 42 ms and the total experiment duration was 1 min (23 frames/sec). The analysis in the presence of specific pharmacological blockers the exposure times were increased to 20 ms, the cycle time was raised to 150 ms to prevent photobleaching (6.6 frames/sec). The total duration time of an experiment depended on the particular applied blocker but normally varied between 5- and 30-min. Pharmacological blockers for SERCA2 [thapsigargin, cyclopiazonic (CPA)], RyR receptor (tetracaine) or gap junctions [carbennoxolone (CBX)] were from Sigma-Aldrich.

### Calcium concentration estimation

Cytosolic calcium was calculated according to the calcium calibration protocol published by Doeller and Wittenberg [44]. Briefly, cultured E10.5 cardiomyocytes were treated first with a calcium free solution and followed by a high calcium solution during imaging recordings. Calcium free or calcium high solutions contained either 15 mM EGTA (Merck) or 25 mM CaCl_2_, diluted both in ACSF supplemented with 10 μM ionomycin. Intracellular calcium concentration was calculated according to the protocol by [44] whereby the K*_D_* value of 225 nM for Fura2-calcium was applied.

### Analysis of cell-cell coupling by Lucifer Yellow

The culture medium of cardiomyocytes was replaced by ACSF and 1% Lucifer Yellow (Invitrogen, L12926) in 90 mM KCl, 3 mM NaCl, 5 mM EGTA, 5 mM HEPES, 5 mM Glucose, 0.5 mM CaCl_2_, 4 mM MgCl_2_) was carefully injected into an individual cardiomoycyte by a glass pipette via the CellTram injector (Eppendorf). After 5 minutes dye spreading to neighboring cardiomyocytes was visualized by Zeiss Axio Examiner A1 using a 40× objective and images were taken using TILL Photonics LA software (version 2.3.0.18, 2013). To measure the area of dye spreading the recorded images from the injected cardiomyocyte and the dye spreading after 5 min were analyzed by the Fiji (ImageJ, version v1.49m, 2015) software.

### Whole-cell patch-clamp recordings of cultured cardiomyocytes

To analyze voltage-gated Na^+^-currents in isolated cardiomyocytes whole-cell patch-clamp recordings were carried out between 3 to 4 days *in vitro*. Cardiomyocytes were visualized under phase contrast optics on an upright microscope (Axioskop, Zeiss) by using a 63×/0.95 water immersion objective. Recordings were performed using a patch-clamp amplifier (EPC-9, HEKA Elektronik). Recording pipettes were filled with an intracellular solution containing (in mM): 10 NaCl, 120 KCl, 5 EGTA, 10 HEPES, 1 MgCl_2_, 1 CaCl_2_, and pH 7.3, 270 mOsmol/kg. The pipette to bath resistance ranged from 3 to 4 MOhm. Series resistance compensation was applied as much as possible (50– 70%). The effective series resistance was in the range of 10–20 MOhm and was tested throughout the whole experiment by using a short depolarizing pulse (10 mV, 20 ms). Recordings were accepted only if the series resistance was less than 20 MOhm. Bath solution contained (in mM): 130 NaCl, 4 KCl, 15 glucose, 10 HEPES, 1 CaCl_2_, and 1 MgCl_2_ (pH 7.3, 310 mOsmol/kg). Whole cell input resistance (R_IN_) was estimated on the basis of passive current responses to moderate depolarizing voltage pulses of short duration (±10 mV for 20 ms). Whole cell membrane capacitance (C_m_) was estimated by integration of the capacitive current transient and division by the respective stimulation voltage. Voltage-activated Na^+^-currents were elicited by a series of 200 ms depolarizing pulses applied from the holding potential of −90 mV, in 10 mV increments between −90 and +50 mV. Passive responses were subtracted by using a hyperpolarizing pulse of −20 mV. Signals were acquired at a rate of 10 kHz and analyzed off-line using WinTida 5.0 (HEKA Electronics).

### Microarray analysis of mRNAs in the embryonic heart

CAR +/+ and CAR -/- E10.5 embryos were dissected and the hearts were flash frozen. RNA was isolated from 3 pooled hearts/sample (n=5; total number of analyzed hearts = 15 for each genotype) by using Qiagen Mini RNA isolation kit accordingly to manufactures’ protocol, followed by Agilent Bioanalyzer quality control. Affymetrix Mouse Gene 1.0 ST and WT PLUS KIT (affymetrix #902464) and GeneChip Hybridization Wash and Stain Kit (#900720) were used as described in the manufactures’ protocol. The microarray was performed in cooperation with the Microarray Facility from the Max-Delbrück-Center. Data sets may be found at the Gene Expression Omnibus (GEO) with accession number GSE138831.

### qPCR to quantify the level of mRNA of selected genes

Isolation of total RNA was performed from freshly dissected E11 hearts according to the instructor’s protocol (RNeasy Mini Kit, Qiagen). cDNA was generated with SuperScript II Reverse Transcriptase (Invitrogen). For the analysis of the expression levels of different genes real-time PCR was performed using SYBR Select Master Mix (Applied Biosystems) as detailed by the supplier. The relative fold change of CAR KO gene expression compared to wild-type hearts was calculated by using the comparative real-time PCR method [45,46]. Actin was used as reference gene. See supplemental information table S2 for primers.

### Biochemical methods

Protein concentrations were determined using the Bradford assay (Bio-Rad #500-0006). Quantification of total protein of connexins in Western blotting was done by solubilizing individual E10.5/11.5 hearts in SDS-PAGE sample buffer and centrifuged. The band intensities were calculated using software Quantity One (Bio-Rad).

### Immunoprecipitation

Immunoprecipitations were done by using covalently labelled IgG fraction of rabbit antibodies to mouse CAR (rb80) to agarose beads by sodium cyanoborohydride and by following the instructions of the Pierce Direct IP Kit (Thermo Scientific, #26148). A HeLa cell line stably expressing connexin43 [47] grown in DMEM/10%FCS/P/S supplemented with 1 μg/ml puromycin (Sigma #P9620) were solubilized in 1% Chaps/TBS/5% glycerol at pH 7.4 supplemented with protease blockers. Insolubilized material was removed by centrifugation (100 000g, 10 minutes). Precipitation was done from 7 (HeLa C×43) mg of solubilized proteins and 20 μg of immobilized anti-CAR IgG.

### Immunocytochemistry and immunohistochemistry

E10.5 embryos were fixed in 4% PFA/PBS for 1.5 h followed by incubation in 15% sucrose (Merck)/PBS for 2 h and overnight incubation in 30% sucrose/PBS. 16 μm thick cryostat sections were incubated in 0.1% Triton X-100/PBS/1% heat-inactivated goat serum using antibodies listed in supplemental table S3. Monoclonal mouse antibodies and corresponding secondary antibodies were incubated on sections using the MOM kit (Vector Laboratories, BMK-2202). Cells were counterstained with nuclear marker DAPI (1 μg/ml) (Sigma). Immuno-stained cryostat sections or cultured cells were analyzed with the LSM 700 confocal microscope (Zeiss, using objectives 10x, 40x, 63x and 100x) and LSM Manager Software (Zeiss) and Fiji/ Image J (Version v1.49m, 2015).

To quantify connexin43 cluster cardiomyocyte cultures were fixed in 4% paraformaldehyde in PBS for 5 minutes followed by solubilization using 0.1% TX-100/0.1% BSA in PBS and washing with PBS/BSA. mAb anti-sarcomeric actinin and rabbit anti-connexin43 and DAPI were applied in PBS/0.1%TX100/5% goat serum overnight. Contacting plasma membranes from two neighboring cells that contained connexin43 spots were encircled and connexin43 spots were counted in confocal images using Fiji software setting the threshold to RenyiEntropy routine; clusters were counted having a size between 0.05 and 1 μm^2^ (2-40 pixels) [48]. The number of spots was related to the encircled area and an average value per cell was calculated. In total 57 wild type and 67 knockout cardiomyocytes were analysed from three embryos for each genotype and roughly 20 spots per cell-cell contact were counted. Then, the average size of connexin43 spots of each cardiomyocyte was calculated. Co-localization of CAR with ZO-1 or connexin43 on plasma membrane regions of cultured cardiomyocytes was analyzed from confocal images by using Fiji/Image J. Co-localizations of CAR with ZO-1 or connexin43 on plasma membranes of cultured cardiomyocytes were analyzed from confocal images by using Fiji/Image J. Calculation of the Pearson correlation was determined by Colo2 derived intensity-based correlation analysis. Costes threshold regression was applied and Pearson coefficient (P) above threshold was used.

For whole mount staining of CAR the IgG fraction of rabbit 80 was applied at a concentration of 2 μg/ml in PBS/0.1%Tx100/5% goat serum for 2 hr at room temperature to paraformaldehyde fixed E10.5 hearts followed by several washing steps and labelling with goat anti-rabbit-Cy3 and DAPI. Hearts were mounted in Immu-Mount (Thermo Scientific, #9990402) for confocal imaging.

### Statistical Analysis

For statistical analysis of data, the SigmaStat software 3.5 (Systat Software, 2006) or GraphPad Prism were used. The Kolmogorov-Smirnov test was carried out to determine whether data are normally distributed. If data sets were normal distributed t-test or paired t-test were applied to measure the significance between the groups of data. The heart organ cultures were analyzed by two-way ANOVA. Other data were analyzed with the Mann-Whitney-Rank-Sum test. Outlier test was performed online using GraphPad QuickCalcs https://www.graphpad.com/quickcalcs/grubbs1. The decay time constants were calculated by a script written in the software IgorPro (WaveMetrics, 2014). Data were represented as means ± SEM. If not given in the Figure legends the following p-values were used to indicate a significant difference between two groups: * p <0.05, ** p <0.01, *** p <0.001.

## 3. Results

### 3.1 The absence of CAR resulted in increased spontaneous beating frequency of cultured embryonic cardiomyocytes

CAR ablation has been shown to disrupt electrical conduction between cardiomyocytes in the adult heart and was correlated with reduced expression of C×45 and C×43 [17,51]. In one of these mutant mouse a raise of the maximal heart rate to 870 in comparison to 730 beats per minute in the wild-type was measured [17]. To analyse the role of CAR in beating of the murine heart at the cellular level, we cultured embryonic cardiomyocytes at high density on fibronectin-coated coverslips. This culture system allowed us to examine cellular calcium cycling, as well the electrophysiological properties of CAR KO cardiomyocytes. Global CAR-deficient mice die between embryonic day 11.5 and 13.5 due to malformation of the embryonic heart [10,13,18]. Therefore, we prepared cardiomyocytes from E10.5 hearts from littermates of wildtype (WT) and CAR-deficient mice. Two days after plating, cardiomyocytes of both genotypes began to beat spontaneously and exhibited synchrony after three days. As indicated by anti-sarcomeric α-actinin staining, CAR was uniformly localized to the surface of cardiomyocytes (Figure 1A). After 4-5 days *in vitro* (DIV), we assessed the beating frequency of the cardiomyocytes using manual counting (Figure 1B) and calcium imaging with the ratiometric calcium indicator Fura2 (Figure 1C and D).

**Figure 1.**
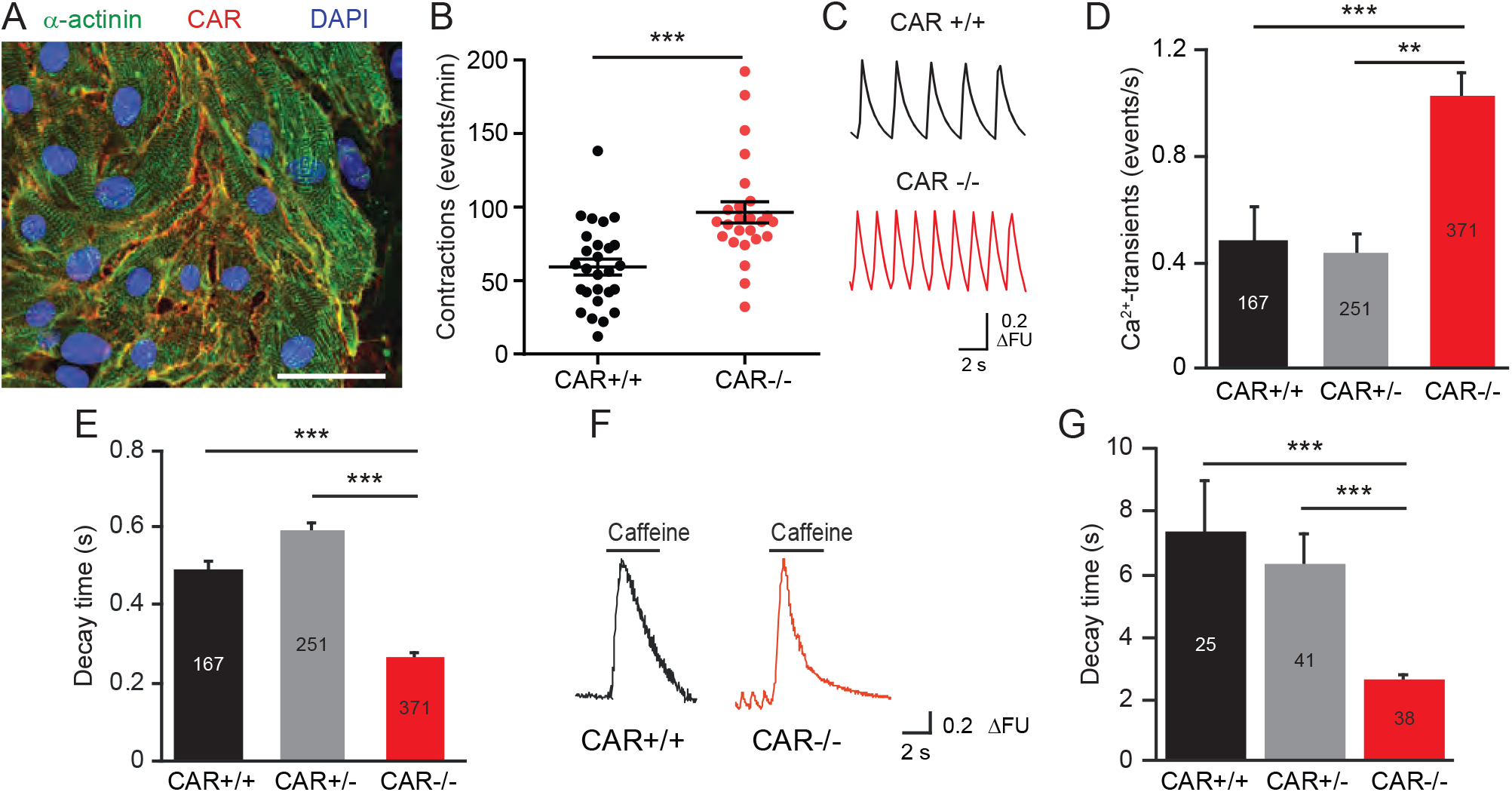
Increased frequency of beating and of calcium transients in monolayer cultures of embryonic cardiomyocytes in the absence of CAR. A) Localization of CAR in cultures from embryonic E10.5 hearts cultured 4 DIV in DMEM/FCS. Cardiomyocytes were identified by labelling with an anti-sarcomeric actinin antibody (green). Scale bar, 80 μm B) The beating frequency of cardiomyocytes was counted manually at DIV 4 at 37 °C in a humidified atmosphere of 95% air and 5% CO_2_ in DMEM/FCS by using an inverted microscope. Cells were from 13 independent cultures and in total 27 wild type and 25 knockout cell clusters were counted. C) Individual traces of calcium transients of cultured cardiomyocytes are shown. FU, relative fluorescence units. D) Summary of the frequency of calcium transients from wild type, heterozygote and CAR-deficient cardiomyocytes. Measurements were done from Fura-2-loaded cardiomyocytes in ACSF at room temperature. Numbers in columns represent numbers of cell clusters from independent cultures. E) Decay time constants of calcium transients of wild type, heterozygote and CAR-deficient cardiomyocytes. F and G) Individual traces of caffeine-induced calcium transients and their decay time constants are shown.

Surprisingly, CAR-deficient cardiomyocytes showed an increased beating frequency (96 ± 7 bpm) compared to WT cardiomyocytes (59 ± 5 bpm, Figure 1B). In line with these measurements, CAR KO cardiomyocytes exhibited significantly increased spontaneous calcium cycling, which correlated with a faster decline of individual calcium transients (Figure 1C and D). The calcium transients for CAR KO cells consistently showed a significantly shorter decay time constant compared to WT calcium transients (Figure 1E). In addition, we also observed a significantly decreased decay time constant in mutant cardiomyocytes after caffeine application, which triggers a complete release of calcium stored in the sarcoplasmic reticulum (Figure 1F and G). Caffeine-induced transients are long-lasting in comparison to spontaneous calcium transients.

These results suggest that the increased beating rate of cultured CAR-deficient embryonic cardiomyocytes could be linked to alterations in calcium cycling, intracellular calcium levels, calcium extrusion from the cytosol, calcium release from internal stores or ionic currents. Alternatively, the increased frequency of calcium transients could result from impaired cell-cell communication. To distinguish between these possibilities, we examined calcium cycling, as well as the electrophysiological properties and cell-cell coupling of WT and CAR-deficient cardiomyocytes.

#### 3.1.1 Mechanisms of calcium cycling were not impaired in CAR-deficient cardiomyocytes

As changes in intracellular calcium levels can influence the beating frequency of cardiomyocytes, we next estimated the cytosolic calcium concentration in WT and CAR-deficient cells, as detailed elsewhere [45]. Both genotypes showed calcium concentrations of 120-130 nM and 350-450 nM during diastolic and systolic phases, respectively, indicating no significant differences in intracellular calcium levels (Figure 2A). In addition, the total calcium content stored in the sarcoplasmic reticulum, which was deduced from the amplitudes of calcium transients after application of 10 mM caffeine, was similar in both genotypes (Figure 2B) [45]. Further, we observed no differences in calcium leakage from the sarcoplasmic reticulum when blocking ryanodine receptors with 1 mM tetracaine (Figure 2C).

**Figure 2.**
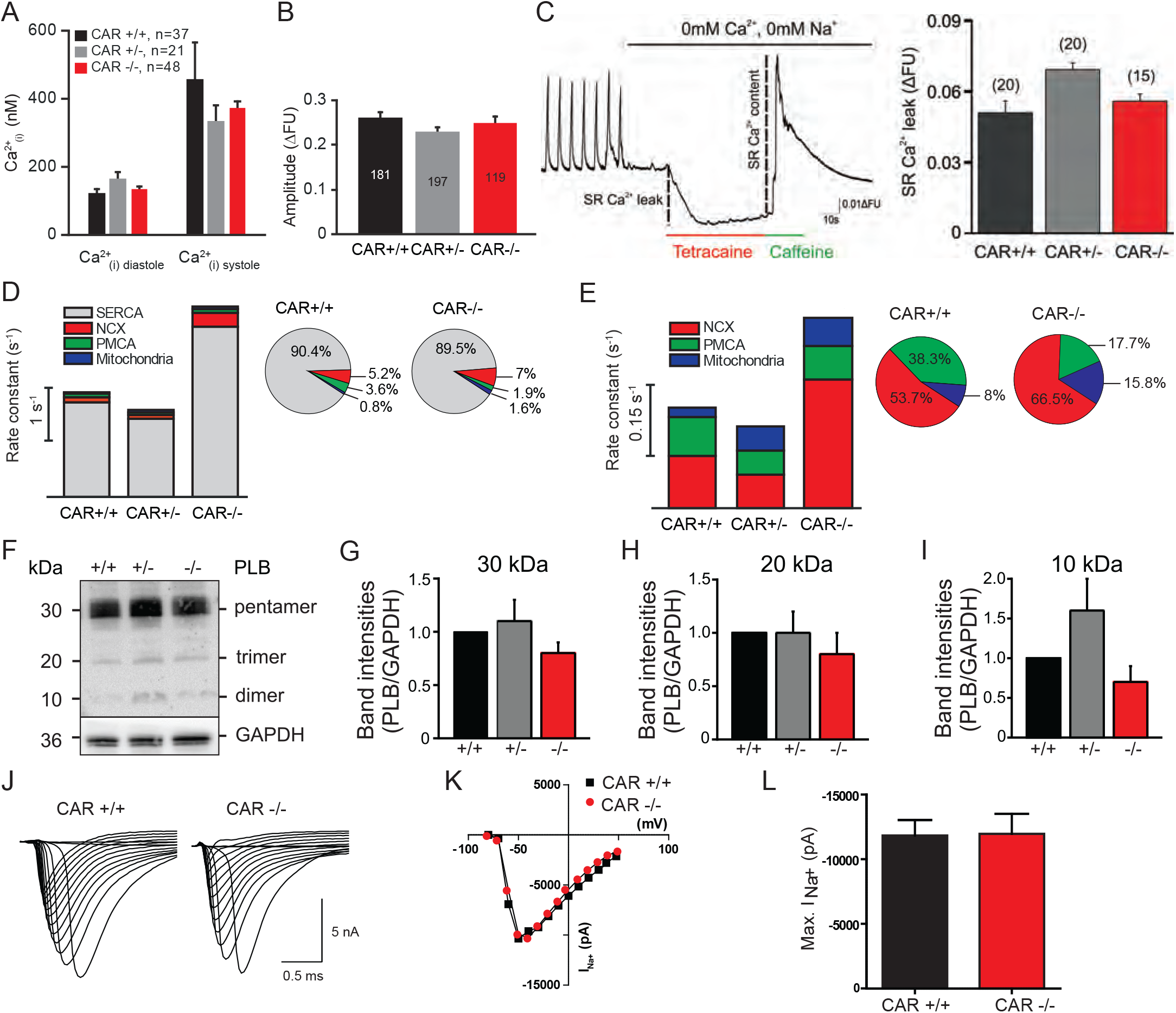
Calcium extrusion is enhanced in CAR knockout cardiomyocytes which correlates with the increased beating frequency. A) Intracellular calcium concentrations at diastolic and systolic phases of wild type, heterozygote and CAR-deficient cultured cardiomyocytes are depicted. B) Amplitudes of calcium transients after application of 10 mM caffeine which might be considered as a measure of calcium levels stored in the sarcoplasmic reticulum. No differences were observed between the different genotypes suggesting that similar amounts of calcium are stored in the sarcoplasmic reticulum. C) Spontaneous calcium events via the RyR, so called calcium-leaks, can induce spontaneous beating and fibrillation of the heart [43,78,79]. The Ca^2+^ leak can be quantified by measuring the changes of cytosolic Ca^2+^ in the presence of 1 mM tetracaine (RyR inhibitor) in Na^+^- and Ca^2+^- free ACSF. Tetracaine completely blocked the RyR and therefore the SR Ca^2+^ leak, consequently the cytosolic Ca^2+^ concentration decreased. Wild-type and CAR knockout cardiomyocytes did not show any significant difference in the intracellular Ca^2+^ drop. As further control at the end of the experiment caffeine was applied to the cells which induced a complete SR release of Ca^2+^. There was no difference of the Ca^2+^ transient amplitude between wild-type and CAR knockout cardiomyocytes. D) Rate constants for all four calcium extrusion components were calculated for spontaneous calcium transients (for calculation see supplemental information). SERCA2 and NCX showed significant increased rate constants in CAR knockout cardiomyocytes. The relative amounts of calcium removed by the different mechanisms are shown for both genotypes in percentages at the right. E) In caffeine-induced calcium transients the NCX rate constants was significantly increased in CAR knockout cardiomyocytes compared to CAR wild types. The relative amounts of calcium removed by the different mechanisms are shown for both genotypes in percentages. F - I) Western blot analysis of the band intensities of phospholamban (PLB) did not show any differences between CAR wild type and knockout E10.5 hearts. Quantification of band intensities of the pentamer (G), trimer (H) and dimer (I) were calculated. Since the regulation of SERCA2 by phospholamban might affect the beating frequency we analysed the polypeptide composition of phospholamban in the absence of CAR. The interaction of phospholamban with SERCA2 negatively regulates the calcium removal from the cytosol. Phosphorylation of phospholamban by PKA results in a release of its monomers from SERCA2 which then reassemble into di-, tri- or pentamers [51,52]. J - L) Sodium currents are not altered in CAR-deficient cardiomyocytes. Single traces (J), the current/voltage relationships (K) and the summary (L) of whole-cell patch clamp recordings of cultured cardiomyocytes are depicted.

Finally, we investigated calcium extrusion mechanisms in cardiomyocytes. Systolic calcium extrusion is mainly carried out by SERCA2 (sarcoplasmic/endoplasmic reticulum calcium ATPase) and NCX (sodium calcium exchanger), with minor contributions from PMCA (plasma membrane calcium ATPase) and mitochondria [52]. To investigate all four calcium extrusion components, we calculated their rate constants as previously described [44] (see supplemental information for detailed description and calculations). The rate constant (*k*) is defined as the amount of calcium per second that is removed from the cytosol, and can be calculated as the reciprocal of the decay time constant (τ). In both WT and mutant embryonic cardiomyocytes, calcium was primarily extruded by SERCA2 (Figure 2D), or by NCX when SERCA2 was inhibited by caffeine (Figure 2E). CAR KO cardiomyocytes, which we previously revealed to beat at a faster rate, showed an increased rate constant for both SERCA2 (grey bar) and NCX (red bar). PMCA and mitochondria played only a minor role in calcium extrusion for both genotypes during spontaneous beating. Further, the relative amount of calcium (shown as %) removed by SERCA2, NCX, PMCA or mitochondria was very similar in both genotypes (Figure 2D and E). In cardiomyocytes of both genotypes, about 90% of the cytosolic calcium was removed by SERCA2. CAR KO cardiomyocytes showed a slight increase in NCX-mediated calcium removal compared to WT cells (5% vs. 7%). Additional physiological studies for NCX using different extracellular sodium concentrations did not reveal any differences between CAR-deficient and WT cardiomyocytes (see supplemental Figure S1).

Interaction of phospholamban, which itself is controlled by phosphorylation, with SERCA2 negatively regulates calcium removal from the cytosol and could therefore affect cardiomyocyte beating frequency [53,54]. However, we observed no differences in the composition of di-, tri- or pentamers between WT and CAR KO embryonic hearts (Figure 2F), indicating no deregulation of SERCA2 by phospholamban. Further, the SERCA2 blockers thapsigargin and CPA only partially reduced beating in CAR KO cardiomyocytes, which might be a reflection of the increased beating activity in these cells (supplemental Figure S2).

#### 3.1.2 Electrophysiological properties were not changed in CAR-deficient cardiomyocytes

To analyse of electrophysiological properties of WT and CAR KO cardiomyocytes, we performed whole-cell patch clamp recordings on cultured cardiomyocytes. We observed no differences in I_NA_ current (Figure 2J – L) between WT and CAR KO cells (WT: −338.2±47.6 pA/pF, n=23 and CAR KO: −384.4.3±50.7 pA/pF, n=47, p=0.5173, mean ± SEM, unpaired t-test). Consistently, the mRNA levels of Nav1.5 (Scn5a) and other sodium channels were unaltered between WT and CAR knockout embryonic hearts. The same held true for potassium channel mRNA transcripts (see database entry GEO GSE138831 and supplemental table 1). Further, the input resistance (WT: 435.1±55.5 MOhm, n=23 and CAR KO: 502.7±67.2 MOhm, n=34, p=0.4492, mean ± SEM) and cell capacitance, which is usually taken as a measure of cell size, were similar in both genotypes (WT: 51.9±10.2 pF, n=23 and CAR KO: 36.5±3.5 pF, n=34, p=0.175, mean ± SEM). These data indicate that the increased beating frequency of CAR KO cardiomyocytes is likely not caused by changes in their electrophysiological properties.

Taken together, we conclude that calcium cycling, the associated extrusion machinery and the electrophysiological properties are not disrupted in the absence of CAR. The increased activity in calcium signaling that we measured in CAR KO cardiomyocytes may instead be a reflection of the increased beating frequency in these cells.

### 3.2 Gap junction-mediated coupling is increased in CAR-deficient cardiomyocytes

The propagation of electrical activity between cardiomyocytes, and consequently the beating of these cells, depends on the expression and regulation of gap junctions [55]. Therefore, we asked whether gap-junction-mediated communication might be impaired in CAR-deficient cardiomyocytes. C×43 and C×45 are the major connexins expressed in embryonic myocytes [56]. In both WT and CAR-deficient cardiomyocytes, C×43 was primarily found at contact sites between neighboring cells (Figure 3A). Surprisingly, these C×43 clusters were significantly larger in CAR KO cells [on average 0.292 μm^2^ and 0.3352 μm^2^ for WT and knockout, respectively; Mann-Whitney U-test (p=0.0007)] (Figure 3B). The relative number of small clusters per membrane area was consistently reduced in CAR KO cardiomyocytes compared to WT cells (Figure 3C). On average, we observed a 40% decrease in total C×43 clusters at cell-cell contact sites in CAR-deficient cardiomyocytes (Figure 3D). The increase in large C×43 clusters in CAR KO cells was not accompanied by altered activity of the kinase Akt, which is known to increase gap junction size by phosphorylating specific serine residues on C×43 (Figure 3E) [57].

**Figure 3.**
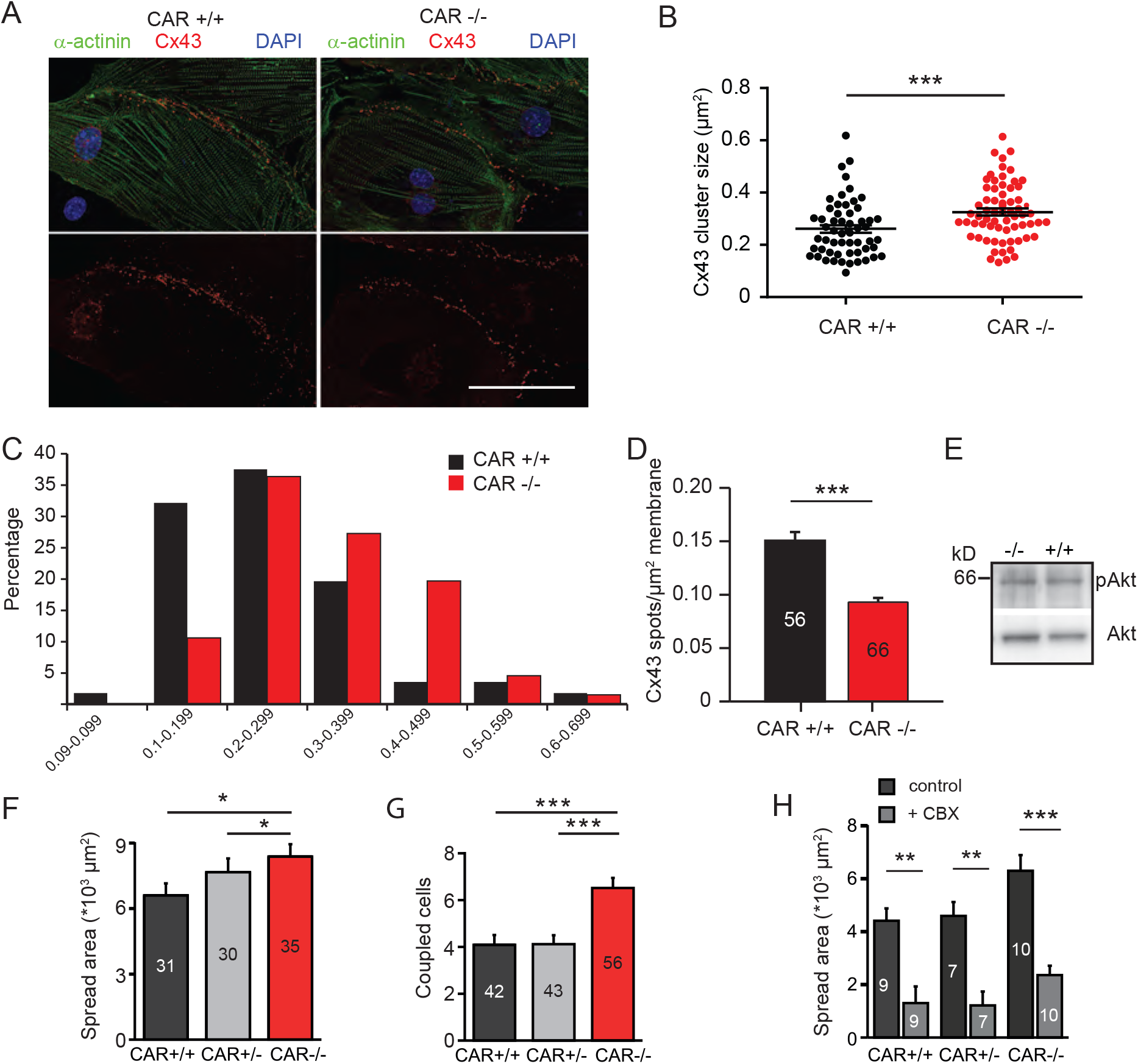
The size of connexin43 spots and dye coupling is increased in the absence of CAR. A-D) Quantification of connexin43 spots at cell-cell contact sites of cultured E10.5 wild type and CAR-deficient cardiomyocytes. Cells were stained by rabbit anti-connexin43, mAb anti-sarcomeric actinin and DAPI. The average size of connexin43 clusters is increased (B, data are from three independent cultures, Mann-Whitney test p=0.0007). In total 56 wild type and 66 knockout cardiomyocytes were analysed. Roughly 20 spots per cell were counted and the average size per cell was calculated. C) The size distribution of connexin43 plaques shows a shift towards larger cluster while smaller ones are reduced (Chi-Square test p=0.0182). D) The total number of connexin43 plaques per cell-cell contact area was reduced. (Mann-Whitney U-test p<0.001). Scale bar in A, 50 μm. E) Western blot of wild type and CAR-deficient embryonic hearts using antibodies to phosphorylated Akt (pAkt) or to total Akt (lower panel) are shown. F and G) Individual cardiomyocyte from E10.5 CAR wild type and CAR knockout cultures were injected with Lucifer Yellow and the spreading was examined after 5 min. The spread area (F) and the number of coupled (Lucifer Yellow stained) cardiomyocytes (G) were significantly increased in CAR knockout cultures. H) Lucifer Yellow spread was blocked by gap junction blocker carbenoxolone (CBX) at 200 μM.

Given the increase in size but decrease in total number of C×43 clusters in CAR KO cardiomyocytes, we next assessed cell-cell coupling using a dye diffusion assay. We microinjected WT and CAR KO cardiomyocytes with Lucifer Yellow, a fluorescent tracer that passes through gap junctions. After 5 minutes, the glass electrode was withdrawn, images were taken and the number of recipient cells (Figure 3G), as well as the spread area (Figure 3F), was measured to quantitatively determine gap junction communication. In comparison to WT cardiomyocytes, CAR KO cells showed a significantly increased number of coupled cells receiving dye from donor cells (Figure 3G), and an increase in dye spread area (Figure 3F). Importantly, coupling was inhibited in all genotypes by the gap junction blocker CBX (200 μM, applied 10 min before Lucifer Yellow application) (Figure 3H).

Further, we observed reductions in C×43 and C×45, but not C×40, C×50 or β-catenin, at the protein level in CAR-deficient hearts by Western blotting (Figure 4A and B). The mRNA levels of C×43 and C×45, as well as a number of other genes, including cell-cell adhesion components and cytoskeletal elements, remained unchanged (Figure 3C and D and table S1, see also database entry GEO GSE138831). qRT-PCR experiments performed for a number of selected genes confirmed the microarray data (supplemental Table S2), indicating that changes occurred at the protein and not at the level of mRNA.

**Figure 4.**
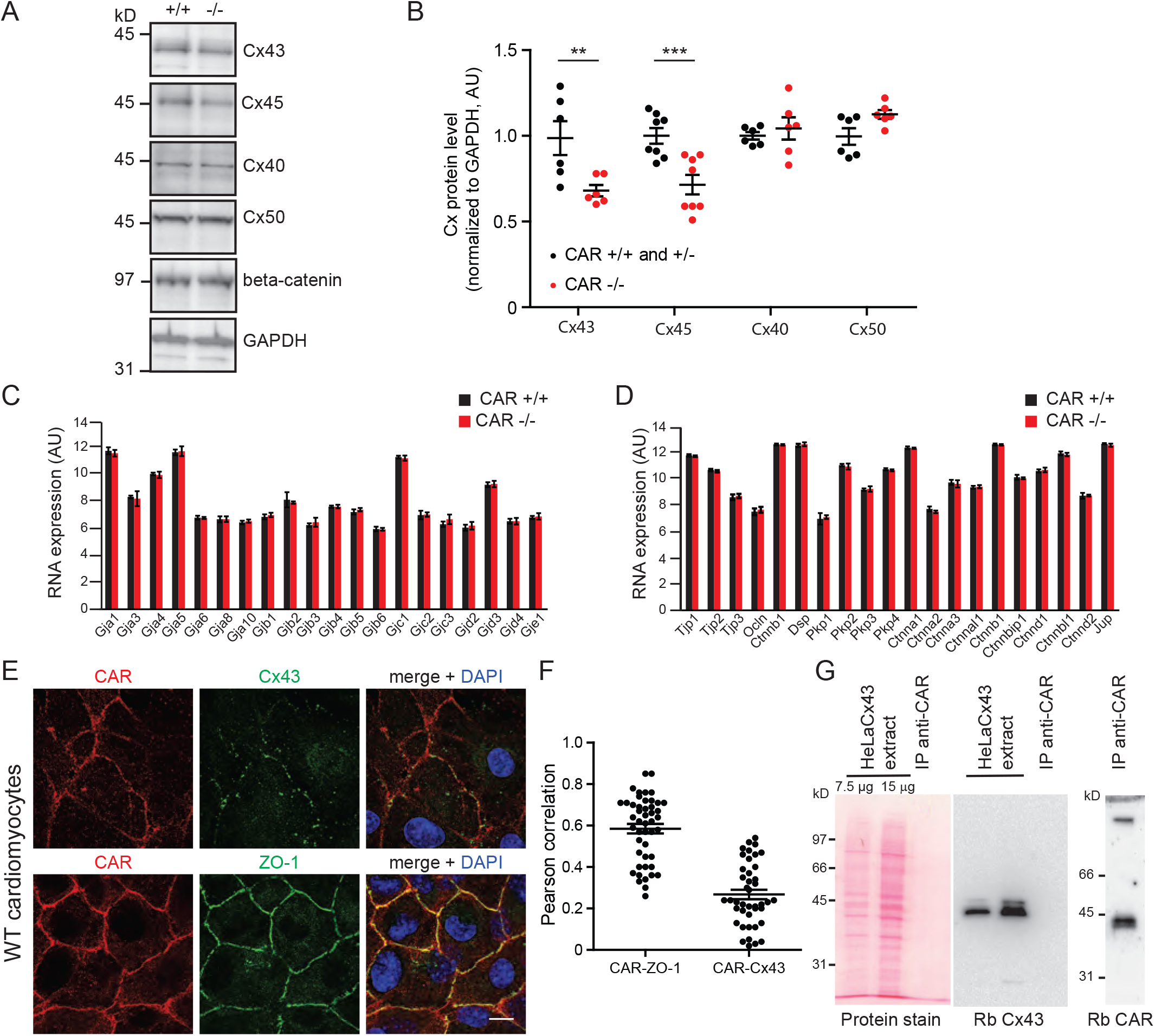
Decreased expression of connexin43 and 45 in CAR-deficient embryonic hearts. A and B) Expression of connexin43, 45, 40 and 50 in wild type and CAR-deficient E10.5/E11.5 hearts shown by Western blots. For comparison β-catenin and GAPDH are shown (from top to bottom). Band intensities from blots were quantified. Reductions were observed for connexin43 and 45 (t-test p=0.0095 and p<0.0001, respectively) but not for connexin 40 or 50 (t-test 0.5773 and 0.1179, respectively) in knockouts. Individual hearts were directly solubilized in SDS-PAGE sample buffer. Molecular mass markers are indicated at the left of the panel. AU, arbitrary units. C-D) Selected Affymetrix microarray data of mRNA expression levels of connexins and cellcell contact proteins of E10.5 hearts of wild type and CAR deficient mice did not reveal differences. For further details see table S1 and data base entry (GEO GSE138831). E-F) On E10.5 cardiomyocytes CAR was not found to co-localize with Connexin43. ZO-1 is shown for comparison. Scale bar, 10 μm. The Pearson correlation of co-localization was calculated using Fiji/Image J software (F). 48 and 43 images were analysed for CAR-ZO-1 and CAR-C×43 co-localization, respectively. G) No co-immunoprecipitation of connexin43 and CAR was detected. CAR was precipitated from extracts of HeLaC×43 by the IgG fraction of rabbit 80 (anti-CAR) directly coupled to CNBr-activated beads. Left panel shows the detergent extract of HeLaC×43 cells stained by Ponceau. (Please note: This protein stain is not sensitive enough to detect proteins obtained by immunoprecipitation). The middle pellet shows the blot with anti C×43 of the HeLaC×43 extract and the IP with anti-CAR. No connexin43 could be detected; however, CAR was easily visualized in the IP (right panel). 7 mg of total protein of the HeLaC×43 extract and 20 μg IgG were used in the IP.

Taken together, these observations indicated a higher degree of cell-cell coupling between CAR-deficient cardiomyocytes compared to WT cells. This suggests that improved electrical propagation via gap junctions could accelerate the beating of CAR-deficient cardiomyocytes. These data are in line with observations that CAR-deficient adult hearts exhibit increased dye coupling between cardiomyocytes and a reduced level of C×43 protein [14,17].

Since CAR appears to affect the oligomerization status of C×43 at the plasma membrane, we investigated a possible direct interaction between CAR and Cx43. In cultured cardiomyocytes, CAR was found in regions of the plasma membrane where Cx43 was not present. Quantification of the co-localization data using Pearson correlation analysis suggested that CAR might not interact directly with Cx43 (Figure 4E and F). Consistently, we found no co-immunoprecipitation of CAR and Cx43 using a HeLa cell line that stably expresses Cx43, suggesting that CAR exert its effect on Cx43 indirectly (Figure 4G). For comparison, the localization of the scaffolding proteins ZO-1 and CAR, which are known to bind to each other [5], are shown (Figure 4E and F).

### 3.3 Increased beating of CAR- deficient embryonic hearts in organ cultures

In the embryonic heart, CAR is expressed on the surface of all cells (Figure 5A and B), but at postnatal stages, becomes predominantly restricted to the intercalated discs (Figure 5B, depicted by arrowheads) [see also for further postnatal stages [26]]. To extend our observations of the increased beating activity of CAR-deficient cardiomyocytes in monolayers, we cultured intact E10.5 hearts from littermates of WT or CAR-deficient mice in Matrigel on chamber slides for up to 72 hours, and the frequency of beating was determined at different time points (Figure 5C). In addition, we applied polyclonal antibodies to the extracellular domain of CAR or “fiber knob” to explant cultures. Both reagents are known to disrupt cell-cell contacts in neurons [2]. “Fiber knob”, here referred to as Ad2, is the tip of the homotrimeric protein of the fiber from the adenovirus capsid which binds CAR on the host cell surface for infection (Figure 5D, E and G). Ad2 binds up to three D1 polypeptides of CAR [58,59] with an affinity higher than CAR to itself, and interferes with cell-cell contact formation of epithelial cells and neurons [2,60]. It interacts specifically with CAR but not with the highly-related proteins CLMP and BT-IgSF (supplemental Figure S3) [3].

**Figure 5.**
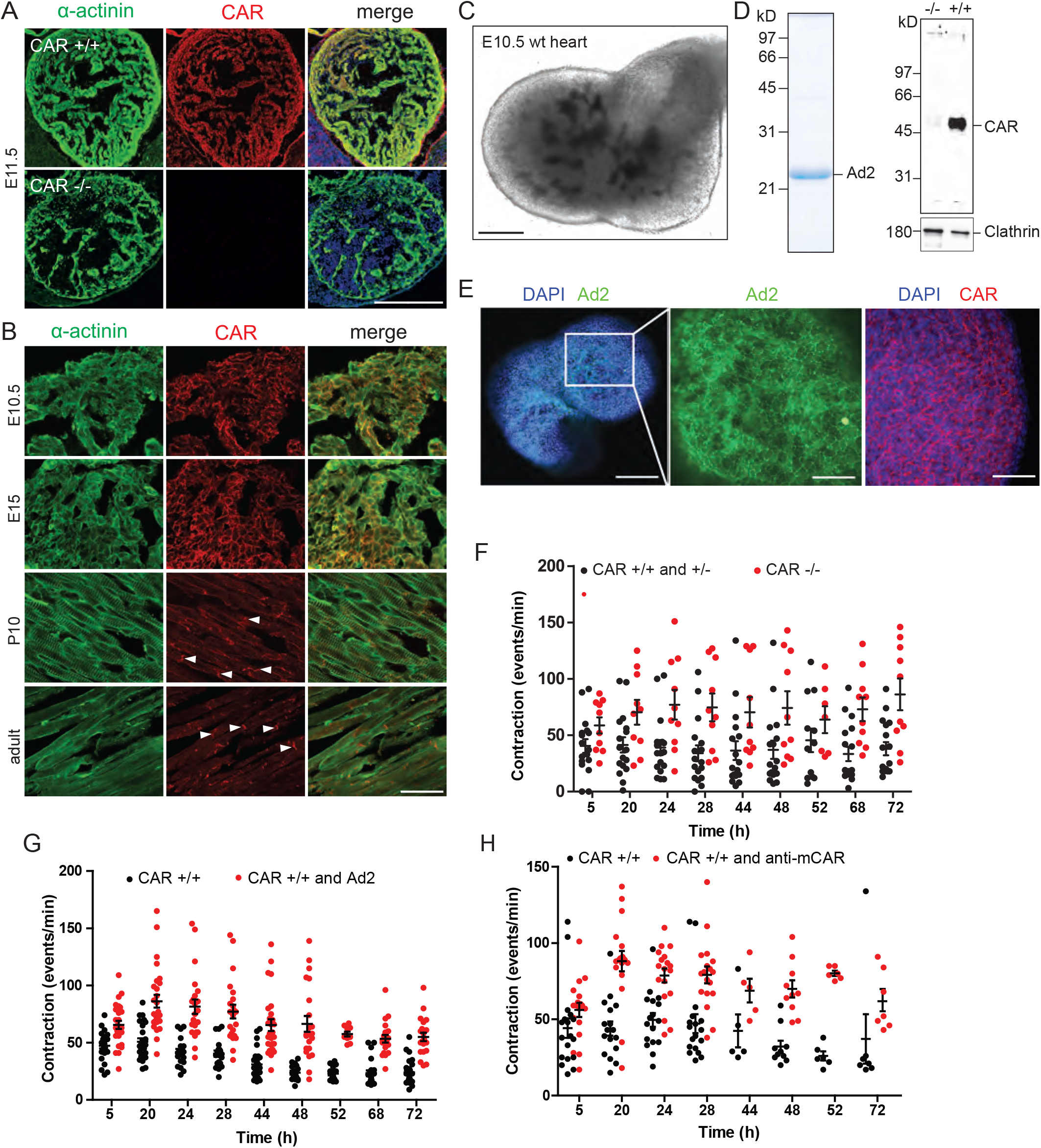
Localization of CAR in the developing and mature heart and increased spontaneous beating frequency of heart organ cultures in the absence of CAR or in the presence of the fiber knob or antibodies to CAR. A) Overview of the localization of CAR in the heart at E11.5. Cryostat sections were stained by rabbit anti-mCAR generated against the extracellular domain of CAR (Rb80) and by monoclonal antibody against sarcomeric α-actinin. Sections from knockout hearts indicated specificity of Rb80. Scale bar, 100 μm. B) Localization of CAR at different developmental stages and in the mature heart. Arrow heads in sections of P10 or adult heart indicate intercalated discs. Scale bar, 50 μm. C) In vitro culture of E10.5 heart in Matrigel to determine the beating rate. Scale bar, 200 μm. D) Purified recombinantly expressed fiber knob Ad2 is shown in 12% SDS-PAGE stained by Coomassie blue. Specificity of the rabbit antibody to the extracellular region of mCAR is demonstrated by Western blotting using wild type and knockout embryonic hearts. Loading was monitored by a monoclonal antibody to clathrin. Specificity of Rb80 is also shown in cryostat section of embryonic hearts in (A). Molecular mass standards are indicated at the left of each panel. E) Whole mounts of E10.5 hearts stained by fiber knob Ad2-Cy2 and Rb80 to illustrate the localization of CAR. Scale bars (from left to right) 190, 60 and 70 μm. F) Spontaneous beating frequency of E10.5 wild type and CAR-deficient heart organ cultures over a period of 72 h. 10 knockouts and 17 wild type/heterozygote hearts were analysed (p<0.0001; two-way ANOVA). G) Spontaneous beating frequency of embryonic wild type heart organ cultures over a period of 72 h in the absence or presence of 0.5 mg/ml fiber knob Ad2. 27 wild type hearts treated with Ad2 and 27 controls were evaluated (p<0.0001; two-way ANOVA). H) Spontaneous beating frequency of embryonic wild type heart organ cultures over a period of 72 h in the absence or presence of 0.5 mg/ml of the IgG fraction of Rb80 to mCAR. 18 wild type hearts treated with Rb80 and 18 hearts as controls were evaluated (p<0.0001; two-way ANOVA).

Regular beating of explanted hearts started after four to five hours *in vitro*. We detected a significant increase in the average spontaneous beating frequency of explanted hearts lacking CAR over several days in culture (Figure 5F). Similarly, the application of Ad2 or polyclonal antibodies to CAR also resulted in an increase in the average spontaneous beating frequency in WT hearts (Figure 5G and H). This suggests that the disruption of CAR’s homophilic binding activity, induced by addition of Ad2 or CAR antibodies, could induce the same effects seen in the CAR KO. Specificity in this culture system was demonstrated by applying antibodies to chick CAR (Rb54), which does not bind to mouse CAR [2] (supplemental Figure S4). Taken together, we conclude that CAR may act as a regulator of gap junctions and heart beating.

## 4. Discussion

The goal of the present study was to examine the function of the Ig cell adhesion molecule CAR in embryonic cardiomyocytes. Unexpectedly, we observed that CAR KO embryonic heart organs and cardiomyocyte cultures exhibit increased spontaneous beating frequency compared to their WT counterparts. This was demonstrated by manual counting and an analysis of calcium cycling, including the mechanisms of calcium extrusion via SERCA2 and NCX. In CAR KO cardiomyocytes, the SERCA2 and NCX rate constants, as well as the proportion of cytosolic calcium removed by SERCA2 and NCX, were significantly increased. However, we observed no changes in the calcium cycling machinery or the electrical properties of these cells. In contrast, the increased beating correlated with increased gap junction dye coupling and an increased size of Cx43 clusters on CAR-deficient cardiomyocytes.

The propagation of electrical activity between cardiomyocytes, which in turn influences the beating of these cells, depends on the regulation of gap junctions [55]. The gating of gap junctions containing Cx43 or Cx45 is primarily regulated by factors such as voltage, pH and phosphorylation [61]. Cell-cell coupling, which is itself modulated by the size and localization of gap junctions, is also crucial for coordinating the spread of action potentials and calcium waves in the heart [62]. Increased cardiomyocyte-cardiomyocyte coupling driven by an increase in junction size could therefore impact the beating frequency [63]. Based on the increased size of Cx43 clusters in CAR KO cardiomyocytes, we hypothesize that electrical excitation pulses in CAR-deficient cardiomyocytes might be more rapidly transferred to neighboring cells, which in turn might result in their faster depolarization. Importantly, our data corroborates the results of a previous study in which CAR-deficient adult hearts showed increased cardiomyocyte coupling and a decrease in Cx43 and Cx45 levels [14,17]. Here the maximal heart rate raised from 730 in the wild type to 870 beats per minute (120%) in the mutant mouse [17]. A role for connexins in regulating cardiomyocyte beating frequency was also shown for Cx45 KO embryonic stem cell-derived cardiomyocytes [64] and in a model with conditional deletion of Cx43 in the mature heart. This resulted in a reduction of the conduction velocity [65,66]. Dysfunction and malformation of the developing heart has also been observed in the absence of Cx43, Cx45 or Cx40, which disrupted gap junction communication [67–71].

The effects of CAR KO shown here are reminiscent of those observed in knockouts of the CAR-related cell adhesion proteins CLMP and BT-IgSF. In mature CLMP-deficient mice, Cx43 and Cx45 levels are reduced in smooth muscle cells of the intestine and ureter, resulting in uncoordinated contractions [50]. In BT-IgSF knockouts, Cx43 is mislocalized in Sertoli cells of the testes and reduced in astrocytes, which impairs cell-cell coupling [72,73]. Together with our observations that CAR plays a role in the adult heart, these data suggest that CAR and the highly-related cell adhesion proteins CLMP and BT-IgSF might regulate the localization and the oligomerization status of Cx43 [4].

An interesting question for future research is the mechanism by which CAR exerts its influence on Cx43. Since connexin mRNA levels were not reduced in the CAR KO, CAR might provide a signal that facilitates Cx43 biosynthesis or might stabilize the expression or oligomerization status of Cx43 within the plasma membrane. However, a direct interaction is unlikely as we observed no co-localization or co-immunoprecipitation of CAR with Cx43. The scaffolding protein ZO-1 is also known to regulate gap junction size [74]. ZO-1 directly interacts with the C-terminal segment of Cx43, which might include its second PDZ domain [75–77]. Inhibition of this binding leads to the uncontrolled formation of large gap junction clusters in culture [74,78,79]. We observed co-localization between CAR and ZO-1 in cultured embryonic cardiomyocytes, which has previously been demonstrated in epithelial cells [5,7]. Although we could not co-precipitate CAR and ZO-1, which might due to the harsh extraction conditions, it is still conceivable that CAR could influence Cx43 gap junction size, in part, via ZO-1. For example, the absence of CAR might cause an inhibition of the binding between Cx43 and ZO-1, which in turn might promote the uncontrolled growth of gap junction clusters.

Interestingly, we observed that reagents that bind to the extracellular region of CAR, such as Ad2, interfered with the beating of embryonic hearts in culture. Crystal structures have been solved for the complete CAR extracellular domain, as well as a complex consisting of the CAR N-terminal-located V-type domain of CAR bound to Ad2 [2,29,30,58]. Conserved amino acid residues within the GFCC’C” surface (650 Å^2^) of the CAR V-type domain are implicated in the homophilic binding of CAR. This area of CAR overlaps with the region that interacts with Ad2 [58]. The stimulatory effect of Ad2 on the beating of heart organ cultures might therefore result from the disruption of CAR-CAR interactions between neighboring cells. Further studies will be required to determine if trans homophilic binding of CAR can modulate the organization of connexins in cardiomyocytes. In summary, we conclude that CAR may regulate gap junctions in murine cardiomyocytes. Candidates that might target CAR in this context include full-length or fragments of AD2, and nanobodies specific to the CAR extracellular domain.

## Acknowledgements

The technical assistance and mouse husbandry of Anne Banerjee, Mechthild Henning, Karola Bach and Petra Stallerow is greatly acknowledged. We are thankful to Dr Marcus Semtner (MDC) for the script written in IgorPro to measure decay time constants, Dr Paul Freimuth (Brookhaven National Laboratory, Upton, USA) for the cDNA encoding the fiber knob Ad2C428N and the Microarray facility of the MDC. We thank Dr Ingo Morano (MDC) for discussion on cardiomyocyte beating. A HeLa cell line stably expressing Cx43 (HeLaCx43) was kindly provided by Prof. Willecke (Molecular Genetics and Cell Biology of Intercellular Gap junctions, Bonn, Germany). FGR is an emeritus professor at the MDC and thanks Dr Carmen Birchmeier (MDC) for generous support and insightful discussions. This work was supported by the MDC and DFG grant SFB 665 (grant B2) to FGR.

## Author contributions

CM (calcium imaging, qRT-PCR, histology, cell culture, microarray, Western blotting, microscopy), RJ (electrophysiology, microscopy), FGR (cell culture, immunocytochemistry, Western blotting, immuno-precipitation, microscopy, biochemistry) and MG (microarray evaluation) performed experiments. CM, MG, RJ and FGR evaluated data. CM, RJ and FGR contributed to the writing of the manuscript.

## Data Availability Statement

Data sets may be found at the Gene Expression Omnibus (GEO) with accession number GSE138831.

## Supplemental Figures and information

**Figure S1**

**Contribution of NCX to calcium extrusion in CAR wild-type and knockout.**

A-B) To study the role of NCX further decay times of caffeine-induced calcium transients were analysed in ACSF without sodium and calcium. Under these experimental conditions NCX is completely blocked. Consequently, decay times of calcium transients increased dramatically for all genotypes. However, no differences between wild type and knockout were observed supporting our conclusion that NCX contributes to the increased extrusion of calcium in CAR knockout.

C-D) The activity of NCX can be manipulated by variations of the driving force, e.g. by changes of the extracellular sodium concentration. Decreasing extracellular sodium concentration from 130 mM to 50 mM caused a strong decrease in the spontaneous beating frequency while an increase in NCX activity by raising the extracellular sodium concentration from 130 mM to 180 mM resulted in increased beating of cardiomyocytes of all genotypes. However, comparison of wild type and knockout always revealed a stronger increase in knockout cardiomyocytes also suggesting a stronger induction of NCX activity in CAR knockout cardiomyocytes.

**Figure S2**

Thapsigargin or CPA only partially reduced the activity of SERCA2 in the absence of CAR.

A and B) Recording of calcium transients during a 20 minutes period showed a strong reduction of the beating frequency of wild type cardiomyocytes after application of 10 μM thapsigargin. In contrast, CAR knockout cardiomyocytes did not change their spontaneous beating frequency indicating that SERCA2 either becomes less sensitive to thapsigargin. In A) individual traces are shown.

C and D) Application of the SERCA2 inhibitor CPA at 10 μM only partially reduced the beating frequency within 20 minutes recording period in CAR-deficient in contrast to wild type cardiomyocytes. In wild type cardiomyocytes amplitudes were already strongly reduced after 1 minute of application of CPA. In C) individual traces and in D) a summary of measurements is shown.

**Figure S3**

**The fiber knob Ad2 binds to the extracellular segment of CAR but not to CLMP or BT-IgSF.**

A) SDS-PAGE of purified Fc fusion proteins analysed by SDS-PAGE.

B) Fc fusion proteins of mCAR, mBT-IgSF, mCLMP or the Fc-fragment at a concentration of 1 μg/ml were immobilised on ELISA plates (200 μl) followed by blocking of residual binding sites and by incubation with 200 μl fiber knob Ad2 at a concentration of 125 ng/ml. Binding was detected by an HRP-conjugated monoclonal antibody directed to the His-tag of Ad2. Binding of Ad2 was only observed to mCAR-Fc but not to BT-IgSF-Fc, CLMP-Fc or to the Fc-fragment (n=3).

**Supplemental Figure S4**

**Analysis of spontaneous heart beating in the presence of rabbit antibodies to chick CAR.**

A-B) Wild type cardiomyocytes were cultured in the presence of the IgG fraction of rabbit antibodies to chick CAR (Rb54, at 0.5 mg/ml) and the spontaneous beating was counted. Rabbit antibodies to chick CAR do not recognize mouse CAR [2]. 12 wild type hearts treated with Rb54 and 11 hearts as controls were evaluated (p=0.199; two-way ANOVA).

### Supplemental information: Quantification of calcium extrusion rate constants

The rate constants k (k = 1/decay time constant) of SERCA2, NCX, PMCA and mitochondria were calculated according to Voigt et al. (2012). The systolic calcium decline is the combined extrusion of calcium by SERCA2, NCX, PMCA and mitochondria. Therefore, the following equation (1) can be defined:

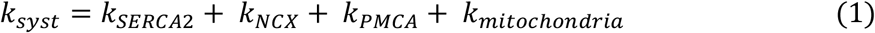

Application of caffeine triggers a complete release of calcium from SR and uptake of cytosolic calcium by SERCA2 is much slower compared to the fast release of calcium. Therefore, during analysis of the calcium decline of caffeine triggered calcium transients SERCA2 is disregarded. The rate constant of caffeine induced calcium transients consequently is composed of the NCX, PMCA and mitochondria extrusion:

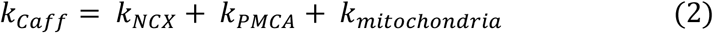

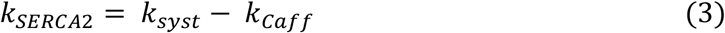

To determine the NCX rate constant Na^+^- and calcium-free ACSF was applied which inhibits NCX activity completely. The cardiomyocytes were treated with 10 mM caffeine and the calcium decline was analyzed. Due to the lack of SERCA2 and NCX activity the calcium extrusion is only performed by PMCA and mitochondria and the following equation (4) can be defined and the rate constant of NCX can be calculated (5):

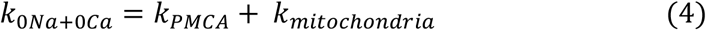

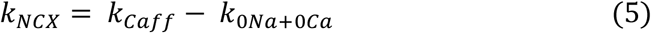

The influence of PMCA on calcium extrusion was determined by blocking NCX activity and mitochondria calcium uptake with 20 μM Ru360. The caffeine induced cytosolic calcium increase is only reduced by PMCA. Therefore, the rate constant of PMCA can be determined by equation (6) and consequently the rate constant of mitochondria by (7):

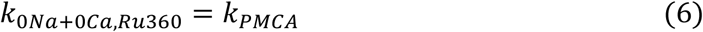

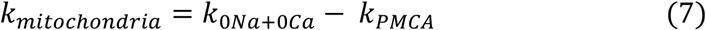

**Supplemental Table S1: Affymetrix gene microarray analysis** (provided as Excel sheet, see also data base entry GEO GSE138831)

**Supplemental Table S2:**
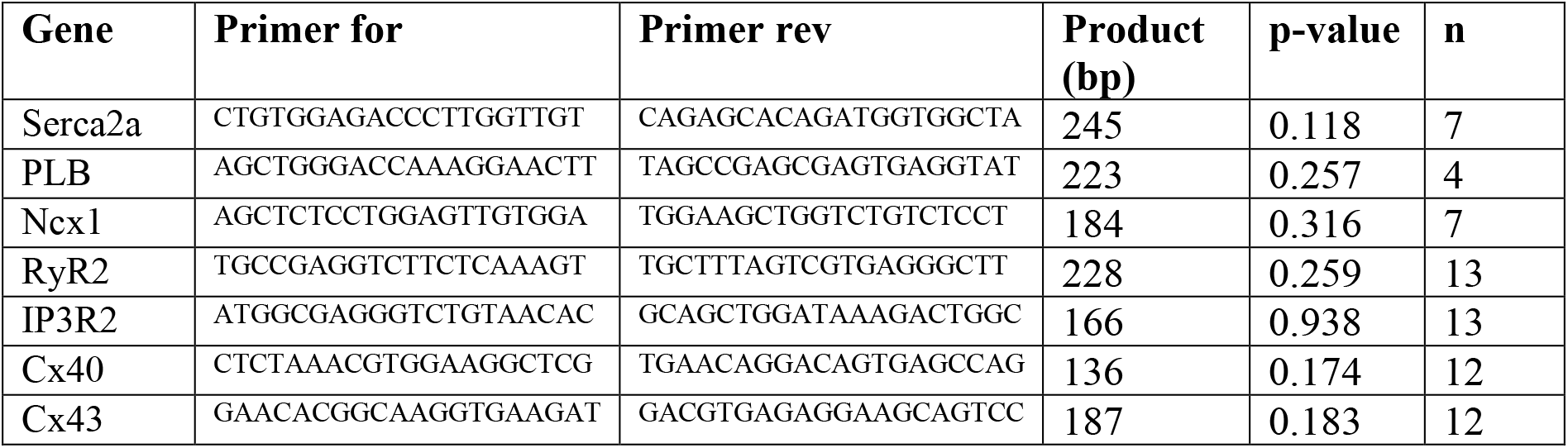
Primer sequences for real-time PCR. To confirm microarray data the expression of several genes was analysed by quantitative RT-PCR and normalized to actin. The data supported the results obtained by Affymetrix microarrays and did not show significant changes of the level of mRNA between wild type and CAR-deficient embryonic hearts for the genes listed below (data not shown).

**Supplemental Table S3:** Antibodies for immunohistochemistry or immunocytochemistry and for Western blotting.

| Antibody | IHC and ICC | Western blots | Source |
| --- | --- | --- | --- |
| Rabbit anti-mCAR-Fc (Rb80) IgG | 1-3 µg/ml | 1 µg/ml | Patzke et al., 2010 |
| Rabbit anti-chCAR-Fc (Rb54) IgG |  | 1 µg/ml | Patzke et al., 2010 |
| Rabbit anti-mL1-CAM |  | 1 µg/ml | Rathjen and Schachner, 1984 |
| mAb anti-Cx43 | 1 µg/ml |  | Transduction Laboratories, #610061 |
| Rabbit anti-Cx43 | 1:200 | 1:1000 | Cell signaling, #3512 |
| mAb anti-Cx40 (B-3) |  | 1 µg/ml | Santa Cruz #sc-365107 |
| mAb anti-Cx45 (G-7) |  | 1 µg/ml | Santa Cruz #sc-374354 |
| mAb anti-Cx50 |  | 1 µg/ml | Invitrogen #33-4300 |
| mAb anti-β-Catenin |  | 0.5 µg/ml | Transduction Laboratories #610153 |
| mAb anti-ZO-1 | 5 µg/ml |  | Invitrogen #339100 |
| Rabbit anti-ZO-1 | 1:100 |  | Invitrogen #40-2200 |
| mAb anti-sacromeric α-actinin (EA-53) | 1:800 |  | Sigma-Aldrich, #A-7811 |
| mAb anti-GAPDH (1D4) |  | 0.5 µg/ml | Novus Biologicals, #NB300-221 |
| mAb anti-Clathrin (heavy chain) |  | 1:1000 | BD Transduction Laboratories #610499 |
| Rabbit anti-phospholamban |  | 1:1000 | Cell Signaling #8495 |
| mAb anti-Penta-His |  | 1:2000 | Qiagen #34660 |
| Goat anti-Rabbit-Cy3 | 1:400 – 1:1000 |  | Dianova |
| Goat anti-Mouse-Alexa488 | 1:400 – 1:1000 |  | Molecular Probes |
| Goat anti-Rabbit-HRP |  | 1:20 000 | Dianova |
| Goat anti-Mouse-HRP |  | 1:20 000 | Dianova |
| Rabbit anti-Akt |  | 1:1000 | Cell Signaling #9272 |
| Rabbit anti-pAkt |  | 1:1000 | Cell Signaling #9271 |

